# Subunit exchange enhances information retention by CaMKII in dendritic spines

**DOI:** 10.1101/372748

**Authors:** Dilawar Singh, Upinder Singh Bhalla

**Author notes:** **Author declaration:** The authors declare no conflict of interest.

## Abstract

Molecular bistables are strong candidates for long-term information storage, for example, in synaptic plasticity. CaMKII is a highly expressed synaptic protein which has been proposed to form a molecular bistable switch capable of maintaining its state for years despite protein turnover and stochastic noise. It has recently been shown that CaMKII holoenzymes exchange subunits among themselves. Here we used computational methods to analyze the effect of subunit exchange on the CaMKII pathway in the presence of diffusion in two different microenvironments, the Post Synaptic Density (PSD) and spine cytosol. We show that in the PSD, subunit exchange leads to coordinated switching and prolongs state stability of the fraction of CaMKII that is present in clusters; and underlies spreading of activation among the remaining CaMKII that is uniformly distributed. Subunit exchange increases the robustness of the CaMKII switch measured as range of bistability both with respect to protein phosphatase 1 (PP1) levels and protein turnover rates. In the phosphatase-rich spine cytosol, subunit exchange leads to slower decay of activity following calcium stimuli. We find that subunit exchange can explain two time-courses of CaMKII activity decay observed in recent experiments monitoring endogenous activity of CaMKII in the spine. Overall, CaMKII exhibits multiple timescales of activity in the synapse and subunit exchange enhances the information retention ability of CaMKII by improving the stability of its switching in the PSD, and by slowing the decay of its activity in the spine cytosol. The existence of diverse timescales in the synapse has important theoretical implications for memory storage in networks.

**Significance Statement:** Despite everyday forgetfulness, we can recall some memories years after they were formed. How are we able to protect some memories for so long? Previous work has shown that the abundant brain protein Calcium/calmodulin dependent protein Kinase II (CaMKII) can form a very stable binary switch which can store information for years. Building on this work, we analyzed the implications of a recently discovered phenomenon of subunit exchange on the state switching properties of CaMKII. In subunit exchange fragments of one CaMKII molecule detatch and exchange with another. We discovered that this improves the information retention ability of CaMKII both in the context where it stores information for long times, and also where it integrates information over the timescale of minutes.

## Introduction

Memories are believed to be stored in synapses, encoded as changes in synaptic strength (1–3). Long Term Potentiation (LTP), an activity dependent change in synaptic strength, is considered to be the primary post-synaptic memory mechanism (4, 5). Various behavioural experiments strongly suggest a critical role for CaMKII in induction of LTP (6, 7). In the CA1 region of Hippocampus, blocking CaMKII activity blocks the induction of LTP (8). After LTP induction, several other pathways including protein synthesis (9), clustering of receptors (10), receptor translocation (11) and PKM-*ζ* activation (12), have been suggested as mechanisms for long-term maintenance of synaptic state. Recent evidence from behavioural assays suggests that CaMKII may also be involved in long-term maintenance of memory (13) (but see (8)).

Any putative molecular mechanism involved in long-term maintenance of memory must be able to maintain its state despite the potent resetting mechanisms of chemical noise and protein turnover. In the small volume of the synapse (~ 0.02 μm^3^ (14)), the number of molecules involved in biochemical processes range from single digits to a few hundred, thereby increasing the effect of chemical noise. Lisman proposed that a kinase and its phosphatase could form a bistable molecular switch able to maintain its state for a very long time despite turnover (15). It has been shown by various mathematical models that CaMKII and its phosphatase PP1 may form a bistable switch (16,17) which can retain its state for years despite stochastic chemical noise and protein turnover (18). Although there is experimental evidence that CaMKII/PP1 is bistable in *in vitro* settings (19, 20), experimental evidence for *in vivo* bistability is lacking. In spine cytosol, CaMKII has been shown not to act like a bistable switch but rather a leaky integrator of calcium activity (8). However, CaMKII may be bistable in special micro-environments such as the “core” PSD where it attaches to NMDA receptor (21, 22).

From computational perspective, the CaMKII/PP1 bistable system is an attractive candidate for memory storage (23). Bistability provides a plausible solution to the problem of state maintenance. Previous modeling work has shown that CaMKII/PP1 system may form a very stable switch despite protein turnover and stochastic noise in the small volume of the synapse (11). The stability increases exponentially with the number of holoenzymes (18). It is important to note that this model exhibits bistable behaviour only in a narrow range of PP1 concentrations in the PSD. This strict restriction may be met because phosphorylated CaMKII is protected from phosphatases in PSD except PP1 (24) which is tightly regulated in the PSD (25).

CaMKII has another remarkable property which was hypothesized by Lisman (26) but discovered only recently, namely, subunit exchange. In this process, two CaMKII holoenzymes can exchange active subunits leading to spread of CaMKII activation (27).

In this paper, we adapt the model of Miller and Zhabotinksy (MZ) (18) to include subunit exchange and diffusion, and quantify the effects of subunit exchange on the properties of the CaMKII-PP1 system in two adjacent neuronal micro-environments: PSD and spine cytosol.

In the PSD, PP1 is tightly regulated and CaMKII is protected from other phosphatases. In the spine cytosol, CaMKII is accessible to other phosphatases along with PP1. We examine how state switching lifetimes in the PSD are affected by subunit exchange in different contexts of PP1 levels, turnover, and clustering of CaMKII. In the spine cytosol we show how the integration of calcium stimuli generates two time-courses of CaMKII activity as a result of subunit exchange (8).

## Results

### Model validation

The basic computational units in our model are individual CaMKII subunits and a CaMKII ring consisting of 6 or 7 CaMKII subunits. We treat the CaMKII ring as a proxy for the CaMKII holoenzyme, which consists of two such rings stacked over each other (28, 29). In our model, CaMKII exists in 15 possible states compared to 2 in (18) (see Materials and Methods). This leads to many more reactions than the MZ model. Since analytical comparison of the two models was not possible, we first compared numerical results from our model without diffusion and without subunit exchange with the MZ model (Fig. 1).

**Fig. 1.**
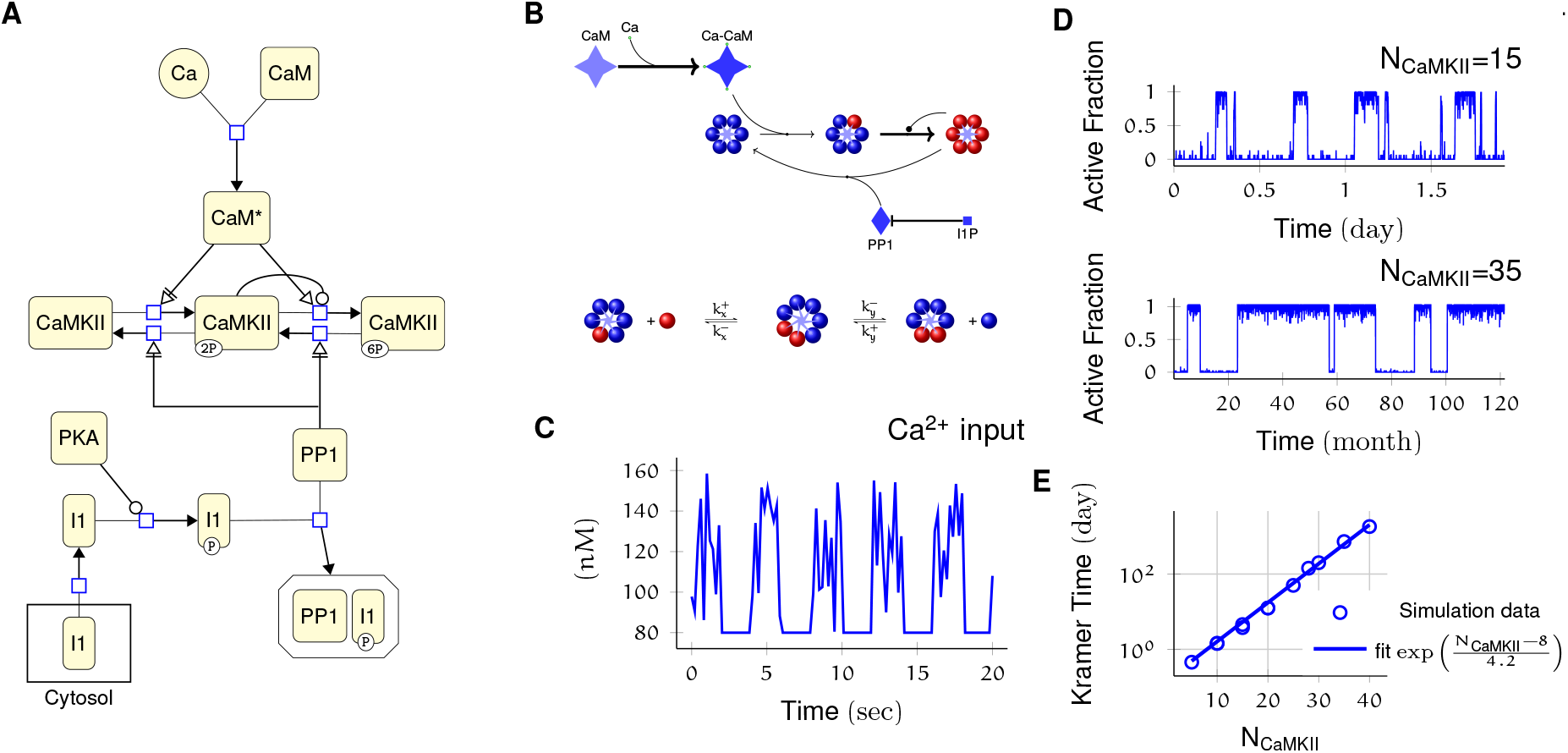
Model description and validation. **(A)** CaMKII/PP1 pathway described in System Biology Graphical Notation (SBGN) - Process Description (PD) Language (30). **(B) (above)** Major chemical reactions in the CaMKII/PP1 pathway. **(below)** Subunit exchange between two CaMKII holoenzymes. Red and blue balls represent phosphorylated and un-phosphorylated subunits respectively. **(C)** Basal calcium (Ca^2+^) profile in spine and PSD in all simulations. Basal Ca^2+^ level is 80nM with fluctuations every 2 s, lasting for 2 s. These fluctuations are sampled from a uniform distribution with mean *μ* =120nM and standard deviation of *σ*=23nM. **(D)** Without diffusion and subunit exchange, CaMKII in our model is bistable. Two trajectories of CaMKII activity (fraction of total CaMKII holoenzymes with at least 2 subunits phosphorylated) are shown for different system size N_CaMKII_=15 **(top)** and N_CaMKII_=35 **(bottom)**. **(D)** Switch stability increases exponentially with system size N_CaMKII_. The average residence time of switch’s stable states increases exponentially with number of CaMKII holoenzymes (N_CaMKII_). Turnover rate v_t_=30h^−1^. Panels **(C, D, E)** show key properties of our model that are very similar to those of the MZ model.

Our model exhibited all the key properties of the MZ model: 1. CaMKII/PP1 under basal calcium stimulus conditions formed a bistable switch in the PSD (Fig. 1C, D), 2. The stability of the switch increased exponentially with system size (Fig. 1E), 3) Increased number of PP1 molecules (N_PP1_) shut off the switch (Fig. 2), and 4. bistability was robust to slow turnover of CaMKII (Fig. 3).

**Fig. 2.**
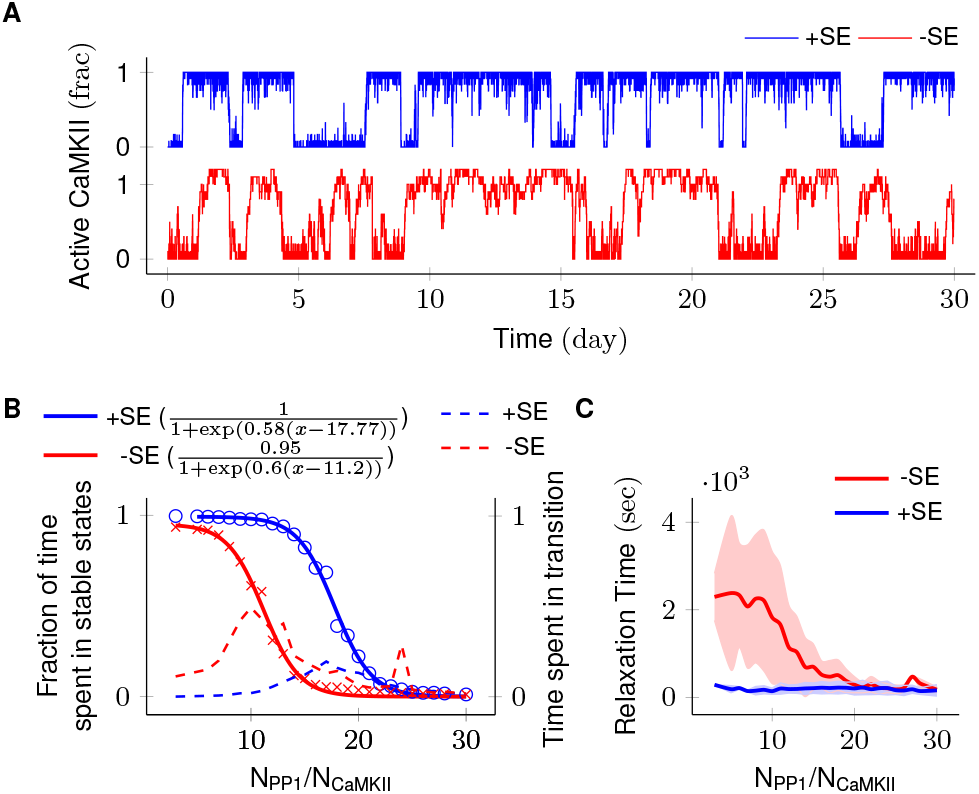
Subunit exchange improves switch tolerance of PP1. **(A)** Two representative trajectories (N_CaMKII_=10) are shown with subunit exchange (_+_SE, blue) and without subunit exchange (_−_SE, red) respectively. **(B)** Blue and red solid-lines represent average activity of switch with and without subunit exchange respectively. The lines are fitted with the function 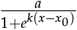. Dotted red and blue lines show the fraction of time that the switch spends in intermediate states (x_a_y_n-a_, 1<a<n-1) with and without subunit exchange respectively. Due to subunit exchange, the switch tolerated a larger amount of PP1 (x_0_ value 11.2 vs 17.77 i.e., a change of 6.57 × N_CaMKII_). Note that the range of PP1 for which switch remains bistable is roughly the same (k, 0.6 v/s 0.58). The fraction of time in intermediate states (dotted lines) is much smaller in the case of subunit exchange (blue dotted line), i.e. the switching time is shorter. **(C)** Due to subunit exchange, relaxation time becomes constant and independent of N_PP1_ (blue vs red). Shaded area represents standard deviation.

**Fig. 3.**
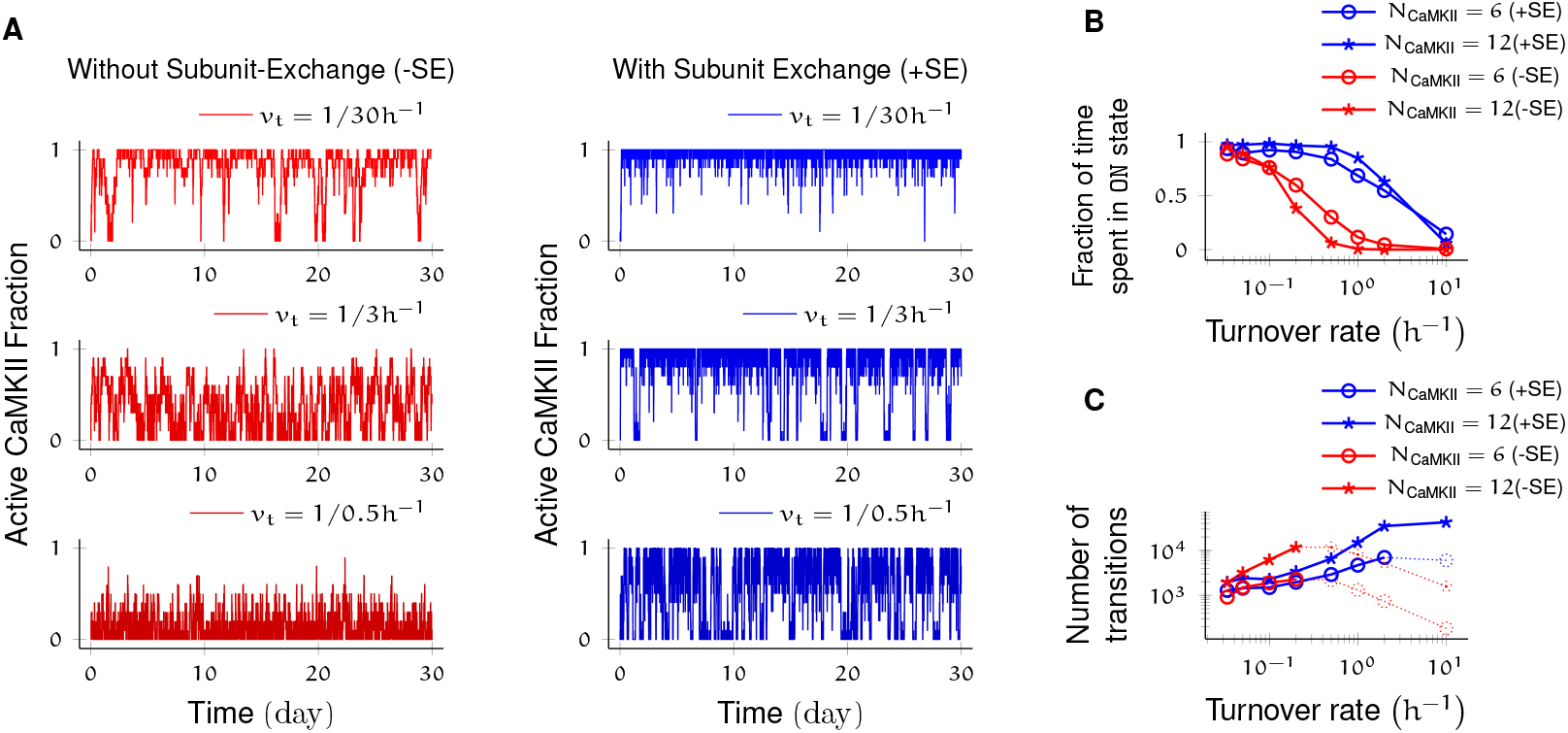
Subunit exchange improves switch tolerance of higher rates of protein turnover. (**A**) Three sample trajectories are shown for a switch of size N_CaMKII_=10 with subunit exchange (_+_SE,blue) and without it (_−_SE,red). We consider three different turnover rates of 1 per 30h, 1 per 3h, and 1 per 0.5 h. As turnover is increased, the state stability of the ON state of the switch decreases. (**B**) Normalized residence time of ON state vs. turnover rate for two switches of size 6 and 12. Without subunit exchange, switch stability decreases exponentially with turnover rate (red), however when subunit exchange is enabled, switch stability is not affected by turnover rates as high as 1 h^−1^ (blue). (**C**) In the bistable regime (solid lines), the number of switching events increases roughly linearly with turnover rate.

Thus, our baseline model exhibited all the key properties that have previously been predicted for the bistable CaMKII switch. However, the subunit exchange and diffusion introduce several interesting additional properties, which we examine now.

### Subunit exchange increases the tolerance of the CaMKII switch to PP1 and to protein turnover

We first analysed switch sensitivity to PP1. In our model as well in the MZ model, the number of PP1 molecules (N_PP1_) had an upper limit for the switch to exhibit bistability. This constraint arises because PP1 must saturate in the ON state of the switch, i.e., the maximal enzymatic turnover of PP1 must be smaller than the rate of activation of CaMKII subunits. However, unlike the MZ model where the addition of one extra PP1 molecule changed the spontaneous state switching time (residence time) of `ON` state by roughly 90% (Fig. 2C in (18)), we did not find lifetime of `ON` and `OFF` states to be very sensitive to PP1. In our model, it required on average 0.5×N_CaMKII_ extra PP1 molecules to cause a 90% change in the residence time. The additional number of PP1 required for switching is roughly equal to number of CaMKII subunits in our model.

We found that the system consisting of N_CaMKII_ holoenzymes remained bistable for N_PP1_=8× to 15×N_CaMKII_ without subunit exchange, and for N_PP1_=12 × to 21 × N_CaMKII_ with subunit exchange. Thus subunit exchange shifted the bistable range to higher values of PP1. Nevertheless, the ratio range in both cases was about the same (blue and red sigmoidal fit in Fig. 2B).

Subunit exchange also had a strong effect on time spent by the switch in transition from one stable state to another (relaxation time). When subunit exchange was enabled, the relaxation time was reduced (red v/s blue dotted line in Fig. 2B) and also became independent of Nppx. Moreover, the standard deviation of the relaxation time was greatly reduced in the presence of subunit exchange (red and blue curve, Fig. 2C).

Parallel results were obtained for the effect of subunit exchange on CaMKII switch robustness in the context of protein turnover. Without subunit exchange, switch stability as measured by residence time of the ON state decreased exponentially with increasing turnover rate. With subunit exchange, however, residence time of ON state remained roughly constant upto a ~10 fold increase in turnover (Fig. 3B), after which subunit exchange could not phosphorylate all inactive holoenzymes produced by turnover, and the switch started to show exponential decay of stability. As expected, turnover increased the number of switching events in the regime of bistability in both cases.

Thus subunit exchange increases the range of N_PP1_ and turnover rate over which the switch remains bistable.

### Subunit exchange facilitates the spread of CaMKII activity

As suggested in (27), we found that subunit exchange facilitates spread of CaMKII activation (Fig. 4). When subunits were allowed to diffuse, they could be picked by neighbouring inactive CaMKII holoenzymes. This effectively overcame the first slow step of CaMKII phosphorylation (Eq. (1)) thereby facilitating the spread of activation.

**Fig. 4.**
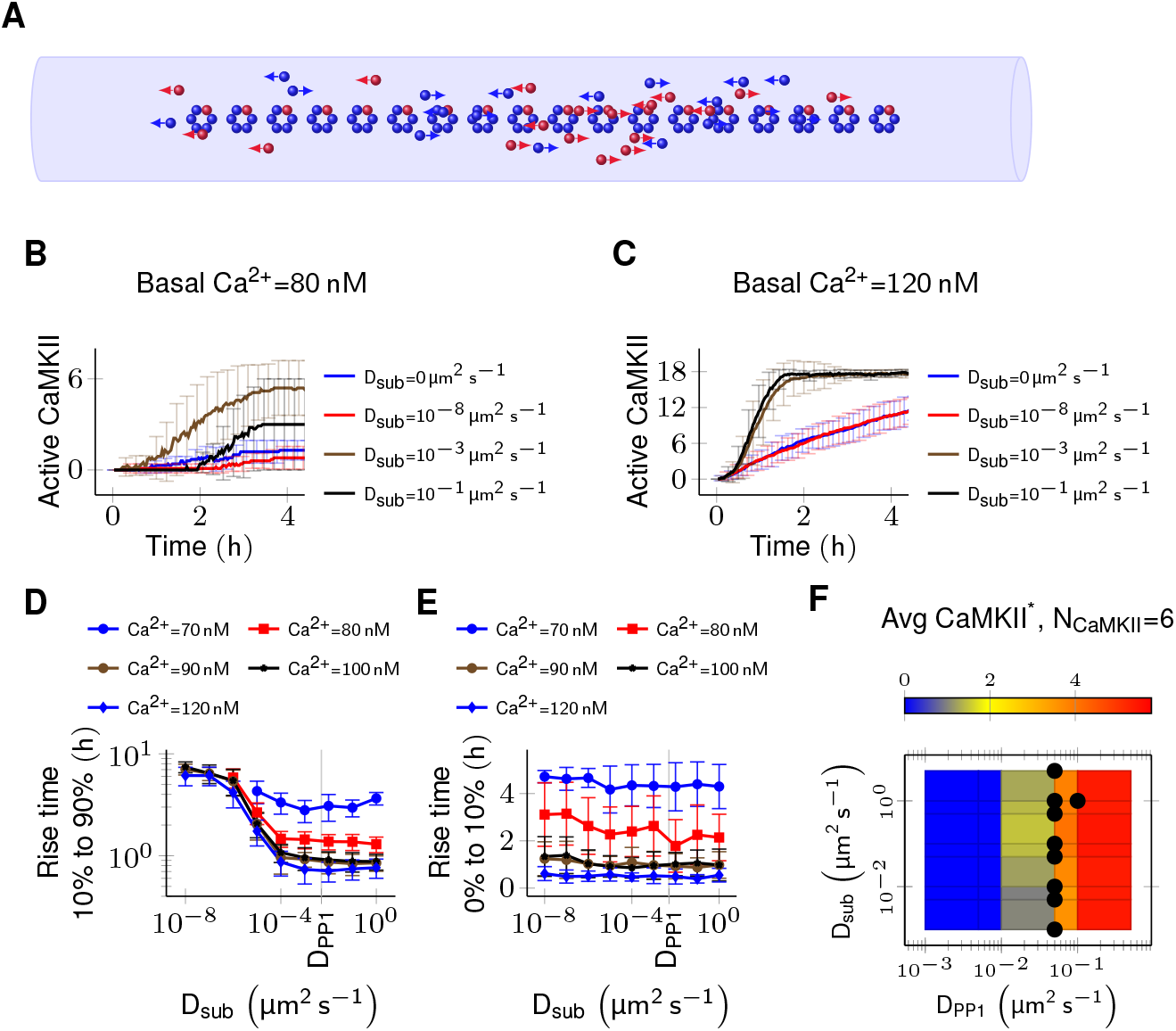
Subunit exchange facilitates the spread of kinase activity (27) but does not change its long-time average. **(A)** 18 CaMKII holoenzymes were put in a cylinder of volume 0.0275 μm^3^ discretized into 18 voxels, each separated by 30ran. For all simulations D_PP1_ = 0.5 μm^2^ s^−1^. **(B)** Activation profile of CaMKII at mean basal calcium level of 80nM (Fig. 1C) for different values of D_sub_. When subunits diffused with zero or negligible coefficient (D_sub_=0 and D_sub_=10~^8^μm^2^ s^−1^), mean activity of CaMKII remained roughly zero. For D_sub_ =0.001 and 0.1 μm^2^ s^−1^, the mean CaMKII activity reached its maximum within 4h. **(C)** CaMKII activates faster at higher mean basal calcium level of 100nM. **(D)** The time taken by CaMKII to rise from 10% to 90% of its maximum value (rise time) in hours v/s D_sub_ for different mean basal calcium levels. The effect of subunit exchange is more prominent at higher calcium levels for all values of D_sub_, and stronger (decreasing rise time) for larger D_sub_ for all values of Ca^2+^ level. 40 trajectories were generated for each trace (Also see Fig. S3). Error bars represents standard deviation. **(E)** The onset of activity time (in hours) v/s D_sub_, where onset of activity time is measured as the time taken by inactive CaMKII to rise from zero to 10% of its maximum value. Average onset of activity time decreased with increasing basal Ca^2+^ level but remained independent of D_sub_. Error bar represents standard deviation. **(F)** Long-time average CaMKII activity (N_CAMKII_=6) is independent of D_sub_, and only depends on D_PP1_. Black dots represent bistable configurations with at least 4 transitions observed in 10 days long simulation.

We put N_CaMKII_=18 inactive holoenzymes in a cylinder with the volume of 0.0275 μm^3^ and the length of 540 nm representing the PSD. The cylinder was divided into 18 voxels (1 holoenzyme in each voxel). Each voxel was separated by 30 nm, which is the average nearest neighbour distance for CaMKII holoenzymes (31). Each voxel was considered to be a *well-mixed* environment i.e. diffusion was instantaneous within the voxel. Diffusion was implemented as cross-voxel “jump” reactions (See Materials and Methods). We did not try 2D/3D diffusion because of its simulation complexity and because it would be expected to be qualitatively similar (32).

We fixed the diffusion coefficient of PP1 (D_PP1_) and quantified the effect of varying the diffusion coefficient of subunits (D_sub_) and basal calcium levels. We used D_PP1_=0.5μm^2^ s^−1^ which is the observed value of the diffusion coefficient of Ras, which is a similar sized protein (33). We ran simulations for 4 hours at basal calcium concentration [Ca^2+^]=80 nM and without subunit exchange (i.e. D_sub_=0). Here the system showed no significant CaMKII activity. When we enabled subunit exchange by setting D_sub_=0.1 μm^2^ s^−1^, Fig. 4B), CaMKII activity rose to maximum within 4 h. As expected, the effect of subunit exchange (rise time quantified as the time taken by CaMKII to rise from 10% to 90%) was stronger when the basal Ca^2+^ was higher (Fig. 4C). Increasing D_sub_ decreased the rise time of CaMKII activity.

Subunit exchange did not have any impact on the average CaMKII activity at longer time scales (Fig. 4E) though we found that long-time average CaMKII activity increased when subunit exchange was enabled. This change was independent of D_sub_ (not due to subunit exchange) but was strongly dependent on D_PP1_. This is due to the fact that potency of PP1 reduced with increased D_PP1_ (Fig. S2) which led to decreased PP1 activity and hence CaMKII activity.

Thus subunit exchange facilitates the spread of kinase activity at short time scale but does not influence its long time activity.

### Subunit exchange synchronizes switching activity of clustered CaMKII

Next we probed the effect of subunit exchange between spatially separated CaMKII clusters at longer timescales. We considered N_CaMKII_ organized into three clusters of N_CaMKII_/3 holoenzymes, each separated by a distance *d*. This configuration corresponds to cases where receptors and CaMKII holoenzymes are clustered at the synapse.

When there is no subunit exchange across voxels i.e. D_sub_=0, these switches are expected to switch independently like multiple coins flipped together, resulting in a binomial distribution of activity. The clustered system had 3 relatively stable bistable systems (long residence time, Fig. 1E). As expected, without subunit exchange, activity in this system had a binomial distribution (Fig. 5B, red plot).

**Fig. 5.**
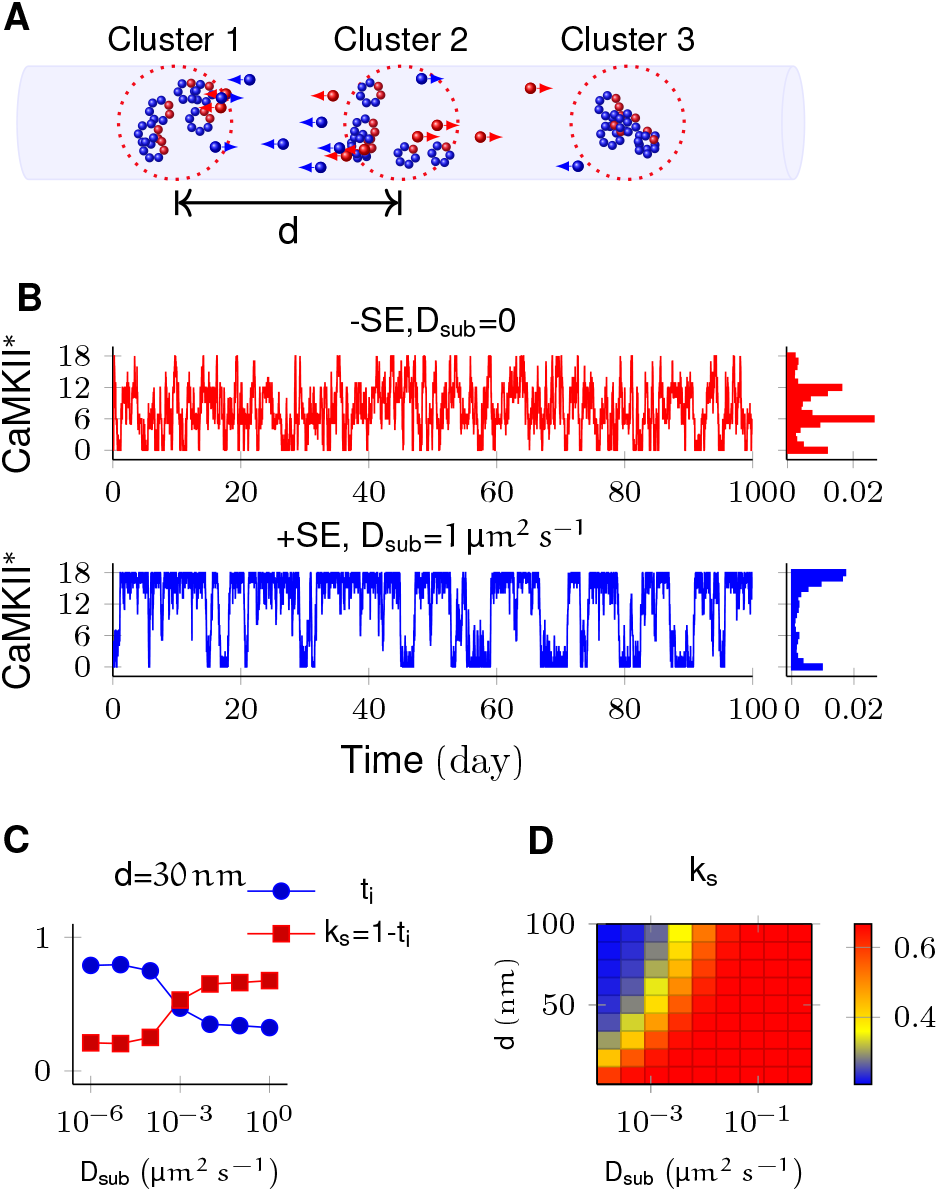
In PSD, subunit exchange synchronizes activity of CaMKII clusters. **(A)** 3 clusters of N_CAMKII_=6 in PSD separated by distance *d*. CaMKII subunits are shown as red (inactive) and blue (active) balls. Subunits and PP1 (not shown) were allowed to diffuse along the axis. **(B)** Without subunit exchange, all three switches flipped independently i.e. state distribution on right is binomial when bin size is N_CAMKII_ (^rΘC^0. With subunit exchange, all switches synchronized their activity i.e. population acted as a single bistable switch (blue). **(C)** Strength of synchronization (k_s_) v/s diffusion coefficient D_sub_ for a system of 3 switches separated from each other by a distance 30 ran. *k_s_* = 1 − *t_i_* where *t_i_* is fraction of time spent by switches in intermediate states x_a_y_n-a_; 1<a<n. Synchronization is strong for k_s_ > 0.4. **(D)** Phase plot of k_s_ v/s D_sub_ and d. The effect of synchronization k_s_ due to subunit exchange is strong and robust to changes in D_sub_, and strong for *d* as large as 100 nm. D_PP1_ =0.5 μm^2^ s^−1^ for all simulations.

Then we allowed PP1 and CaMKII subunits to undergo linear diffusion. We fixed D_PP1_=0.5μm^2^ s^−1^ as before and varied *D*_sub_ to quantify effect of subunit exchange. Subunit exchange led to synchronization of switching activity. The population of clustered CaMKII acted as a single bistable switch (Fig. 5B, blue plot). This effect was strong and robust to variation in D_sub_. Even for a very small value of D_sub_=0.01 μm^2^ s^−1^, we observed strong synchronization (Fig. 5D). The synchronization disappeared completely for diffusion coefficient less than D_sub_=10^−4^μm^2^s^−1^ for distance *d* > 30 nm (Fig. 5D).

Thus for most physiologically plausible values of diffusion coefficient D_sub_, subunit exchange causes synchronization of switching activity of clustered CaMKII.

### Subunit exchange may account for the observed dual decay rate of CaMKII phosphorylation

Finally, we asked if subunit exchange might account for the complex time-course of CaMKII dynamics in spine as observed in recent experiments (8). We designed an experiment to replicate an experiment where CaMKII was inhibited by a genetically encoded photoactivable inhibitory peptide after activating it by glutamate uncaging (34). In the spine, CaMKII is more accessible to phosphatases than in the PSD, where our previous calculations had been located. To model the increased availability of phosphatases, we increased the number of PP1 by an order of magnitude, and increased the volume of the compartment to match the volume of a typical spine head i.e. 0.02 μm^3^ (14). We found that CaMKII acted as a integrator of calcium activity with typical exponential decay dynamics (Fig. 6A). We then enabled the diffusion of CaMKII subunits and PP1 with same diffusion coefficient D_sub_=D_PP1_=1 μm^2^ s^−1^. These conditions decreased the rate of dephosphorylation of CaMKII holoenzymes significantly. The decay dynamics could not be fitted using a simple exponential (Fig. 6B). With our values of parameters, decay rate decreased by an order of magnitude when subunit exchange was enabled (Fig. 6B).

**Fig. 6.**
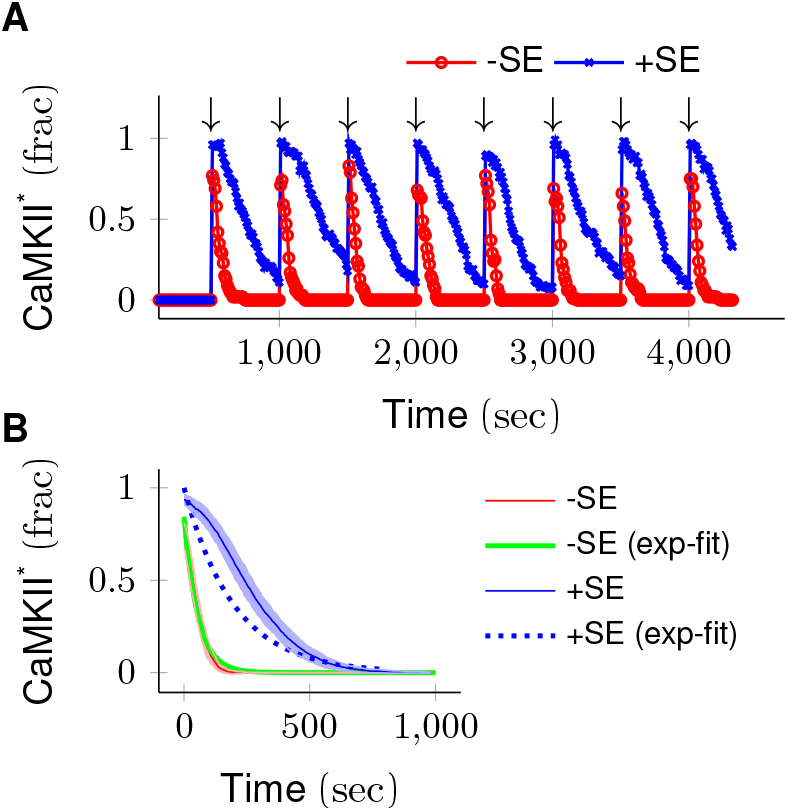
CaMKII acts as a integrator of calcium activity in spine cytosol. Due to subunit exchange, CaMKII decayed with much larger time-constant and almost linearly. **(A)** Trajectories of CaMKII activity (fraction of total CaMKII) are shown in red (without subunit exchange) and blue (with subunit exchange). Every 500s, a 3s long strong calcium pulse is applied to the system (↓) thereby activating CaMKII. After the pulse, calcium levels were brought down to basal level. **(B)** Average decay dynamics for 500s after the onset of strong calcium pulse (↓). Solid red line shows decay dynamics when there is no subunit exchange; CaMKII decays at timescale of approximately 50 s. Solid blue line shows the case when subunit exchange is enabled, CaMKII decay has larger time-constant of ~ 300 s. Dotted blue line shows exponential fit of slow time-course which does not fit well. On the other hand, the fast trajectory fits very well by an exponential (red, `-SE (exp-fit)`). Shaded areas are the standard deviation. Though the absolution value of fast and slow timescales are much higher, the ratio of fast and slow time-courses matches well with experimental data (8).

We expected subunit exchange to have a strong effect on the decay activity of clustered CaMKII in spine cytosol (e.g. CaMKII bound to actin) because of the close proximity of holoenzymes, leading to rapid exchange. Our simulations supported this prediction. If there are populations of clustered as well as non-clustered CaMKII in the spine, we expect that they will exhibit long and short time-courses of activity decay.

Thus we suggest that subunit exchange may be a mechanism that leads to CaMKII it activity decaying with two time-courses in spine cytosol (8).

## Discussion

Here we have shown that subunit exchange strongly affects the properties of CaMKII/PP1 pathway, both in its role as a bistable switch in PSD and as an integrator of calcium activity in spine cytosol. In the PSD, where the model was tuned to elicit bistable dynamics from clustered CaMKII, subunit exchange improved the stability of CaMKII/PP1 switch by synchronizing the kinase activity across PSD (Fig. 6). It also improved CaMKII tolerance of PP1 and turnover rate (Fig. 2 and Fig. 3). In the case where CaMKII was uniformly distributed in PSD, subunit exchange facilitated more rapid activation of CaMKII (Fig. 4BCD) (27). These simulation results predict that a CaMKII mutant lacking subunit exchange would be deficient in the switch stability and slower to activate.

In the spine head, subunit exchange facilitated integration by prolonging the decay time-course of kinase activity (Fig. 6). The fact that CaMKII dynamics changed from an integrator to bistable switch as we moved from spine cytosol (a phosphatase rich environment) to the PSD (where PP1 is tightly controlled) suggests an interesting sub-compartmentalization of functions in these microdomains. Furthermore, we observed that the clustering of CaMKII had important implications for its sustained activity.

CaMKII is non uniformly distributed in PSD. Most of it is concentrated in a small region of 16 nm to 36 nm below synaptic cleft (22) where it may exist in large clusters given that CaMKII has multiple binding partners in the PSD. Our study predicts that subunit exchange may lead to synchronization when CaMKII is clustered, or more rapid activation by calcium when it is uniformly distributed. Given that CaMKII can form clusters with N-methyl-D-asparate (NMDA) receptors, it would be interesting to study the mixed case where some CaMKII is clustered and rest is uniformly distributed. This would require detailed 3D simulation and is beyond the scope of this study.

Subunit exchange is unlikely to have any impact on neighbouring spines. The mean escape time of a single CaMKII subunit from a typical spine is between 8 s to 33 s (35). In a real synapse, this time would be even larger given that CaMKII interacts with many other molecules. Any phosphorylated subunit is almost certain to be de-phosphorylated by PP1 during this time. We therefore predict that the effects of synchronization are local to each PSD, where PP1 is known to be tightly controlled. Subunit exchange loses its potency in the phosphatase rich region of the bulk spine head or dendrite. We therefore consider it unlikely that CaMKII subunit exchange plays any role in intra-spine information exchange such as synaptic tagging.

Finally, we suggest that existence of diverse time-scales of CaMKII activity in the PSD and spine head has important theoretical implications. A very plastic synapse is good at registering activity dependent changes (learning) but bad at retaining old memories. On the other hand, a rigid synapse is good at retaining old memories but is not efficient for learning. A theoretical meta-model which sought to strike a balance between these two competing demands requires that diversity of timescales should exist at the synapse (36). In this model, complex synapses with state variables with diverse time-scales are shown to form a memory network in which storage capacity scales linearly with number of synapses, and memory decay follows 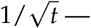 a power-law supported by psychological studies (37). This model requires memory trace to be first stored in a fast variable and then progressively and efficiently transferred to slower variables. Our study suggests a concrete mechanism for such a process. Here, calcium concentration in PSD can be mapped to the fastest variable. The CaMKII integrator in cytosol could represent the second slower variable to which the trace is transferred from calcium. Further, the state information is transferred to the third slower CaMKII bistable switch. The dynamics of CaMKII in the PSD forms an even slower bistable variable for longer retention of the memory trace. It is possible that memory is transferred from here to even slower variables, such as sustained receptor insertion (11), PKM-*ζ* activation (12), or local protein synthesis (9).

## Materials and Methods

The kinase CaMKII has a rich history of modeling spanning over two decades with varying complexity (16). We based our study on the work of Miller and Zhabotinksy (MZ model) (18). We extended their model to incorporate *subunit exchange* and diffusion. We treat the CaMKII ring as proxy for the holoenzyme because vertical dimers (one subunit from the top ring and one from the ring below) are inserted or released together (38) and we assumed that the top and the bottom subunits of a vertical dimer phosphorylate and de-phosphorylate together.

In our model, a CaMKII ring with *n* subunits (*n*=6 or 7) can exist in *n* different states *x_a_y_n−a_* where *a* is the number of un-phosphorylated subunits (represented by *x*), and *n* – *a* is the number of phosphorylated subunits (represented by *y*). All CaMKII rings in our model have either 6 or 7 subunits. The number of CaMKII states in our model increased from 2 (`ACTIVE` and `INACTIVE`) in MZ model to 13. We ignore all rotational permutations and kinetically unlikely cases where there are discontiguous phosphorylated subunits in the ring. We assumed that the phosphorylation of neighbouring subunit proceeds clockwise.

### Phosphorylation and dephosphorylation of CaMKII ring

The activation of CaMKII follows the same dynamics as in MZ model. Upon its influx into spine, Ca^2+^ binds to calmodulin (CaM) to form calcium/calmodulin complex (Ca^2+^/CaM). The first step in CaMKII activation requires simultaneous binding of two Ca^2+^/CaM to the two adjacent subunits of CaMKII. The probability of such simultaneous binding of two Ca^2+^/CaM is very low at basal Ca^2+^ concentration. Once a subunit is phosphorylated, it catalyzes phosphorylation of it’s neighbour (*auto-phosphorylation*). Auto-phosphorylation requires binding of only one Ca^2+^ /CaM therefore proceeds at much faster rate compared to the first step. Once fully phosphorylated, CaMKII moves to PSD where it binds to NMDA receptor. Upon binding, it is no longer accessible other phosphatases save PP1.

Following Zhabotinksy, we also assumed that Ca^2+^/CaM binding to CaMKII ring is independent of the current state of the ring. The activation of a subunit by Ca^2+^ follows a Hill equation (Eq. (1)) (17).

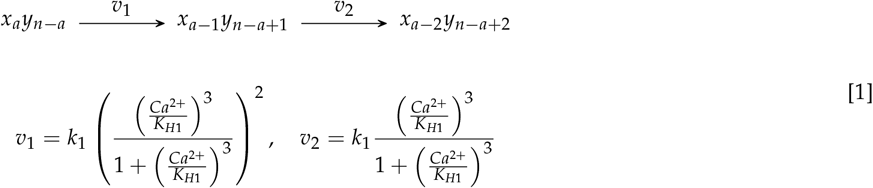

where *n* = 6 or 7, and 1 ≤ *a* ≤ *n*.

The dephosphorylation of the CaMKII ring and the single subunit follow a Michaelis-Menten like scheme. But rather than using the Michaelis-Menten approximation, we implemented this as coupled mass-action chemical reactions as shown by Eq. (2).

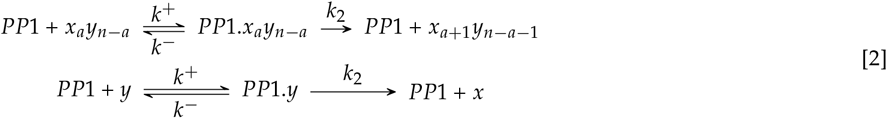

where *n* = 6 or 7, and 1 ≤ *a* ≤ *n*.

Following Miller et al. (18), we assumed *k*^−^ = 0. This gave us 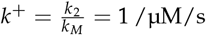.

### Subunit exchange

In our model, a CaMKII ring is made up of either 6 or 7 subunits. Therefore CaMKII ring with 6 subunits cannot lose a subunit while a CaMKII ring with 7 subunits cannot gain a subunit. The gained or lost subunit can be either phosphorylated (*x*) or un-phosphorylated (*y*). All possible reactions which result in either gain or lose of a subunit are given by Eq. (3).

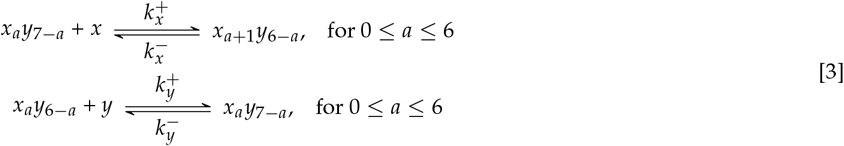

The values of 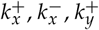, and 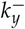 are not available in the literature that we are aware of. We assumed that these reactions operate at the timescale of second. Bhattacharya et. al. (38) speculate that upon activation, the hub of holoenzyme becomes less stable and more likely to open up and lose a subunit. Therefore we assumed the rate of losing subunit 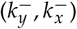 to be larger than the rate of gaining a subunit 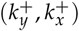. In all simulations, we maintained the following ratio 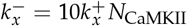 and 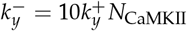.

### PP1 deactivation

In the PSD, PP1 is the primary – and perhaps only – phosphatase known to dephosphorylate CaMKII (39). In the PSD, PP1 is inhibited by phosphorylated inhibitor-1 (I1P) and a dopamine- and cyclic-AMP regulated neuronal phosphoprotein (DARPP-32) (an isomer of inhibitor-1 (I1)) (40, 41) (also see (42)).

I1 is phosphorylated by protein kinase A (PKA) and dephosphorylated by calcineurin (CaN). We followed Zhabotinksy’s assumption that I1 level are constant in the PSD because I1 exchanges rapidly with spine cytosol (17). We assumed the concentration of I1 to be the same in both PSD and spine cytosol. We followed Miller and Zhabotinksy in making the approximation that I1 phosphorylation is very fast compared to the phosphorylation of CaMKII. Therefore, under the *quasi-equilibrium* approximation, I1P level are given by Eq. (4) where *v_PKA_* is the activity of PKA divided by its Michaelis constant, and *v_CaN_* is the activity of CaN divided by its Michaelis constant (18).

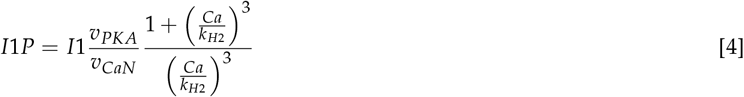

I1P renders PP1 inactive by forming I1P-PP1 complex (I1P.PP1). This reaction (5) is also assumed to be fast (18).

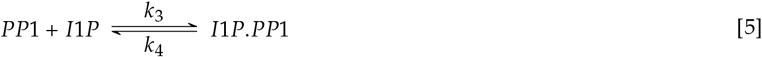

### Turnover

The turnover of CaMKII is a continuous process with rate *v_t_* s^−1^. We modeled turnover by replacing a CaMKII ring with *a* > 1 phosphorylated subunits by an inactive CaMKII ring of the same symmetry.

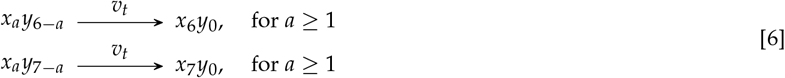

### Diffusion and simulation method

Diffusion is implemented as cross voxel “jump” reaction. Diffusion of a species X with diffusion-coefficient D_X_ between voxel A and B separated by distance *h* is modelled by reaction 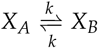 where *k* = *D_X_*/*h*^2^, and [*X_A_*] = [*X_B_*] = [*X*] /2 (43). Lowering *h* improves the accuracy of diffusive component of system but bimolecular reactions are increasingly lost as *h* gets smaller and smaller (44). We have many bimolecular reactions with diffusing species (PP1 and subunits) as reactants. For all of our simulations, *h* is 30 nm – mean nearest-neighbour distance for CaMKII. We computed the critical value of *h* namely *h_crit_* for which error is in acceptable bound i.e. < 1%. The value of *h_crit_* is determined by the fastest bimolecular reaction (Eq. (2)) and the slowest diffusion coefficient. The lower bound on *h* i.e. 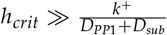, where *k* is the reaction rate (44). Based on our own numerical results (see SI) and other studies (44, 45), we are confident that 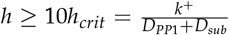 is a good value (45). We have *h_crit_* ≤ 3.2 nm whenever *D*_*PP*1_ + *D_sub_* ≥ 0.5 μm^2^ s^−1^. For all simulations presented in main text, we maintain *h* ≥ *h_crit_*. For a case where *h* is smaller than *h_crit_* in some trajectories see Fig. S2.

All simulations were performed using Stochastic solver (Gillespie method) available in MOOSE simulator (https://moose.ncbs.res.in, version 3.1.2) (46).

**Table 1.**
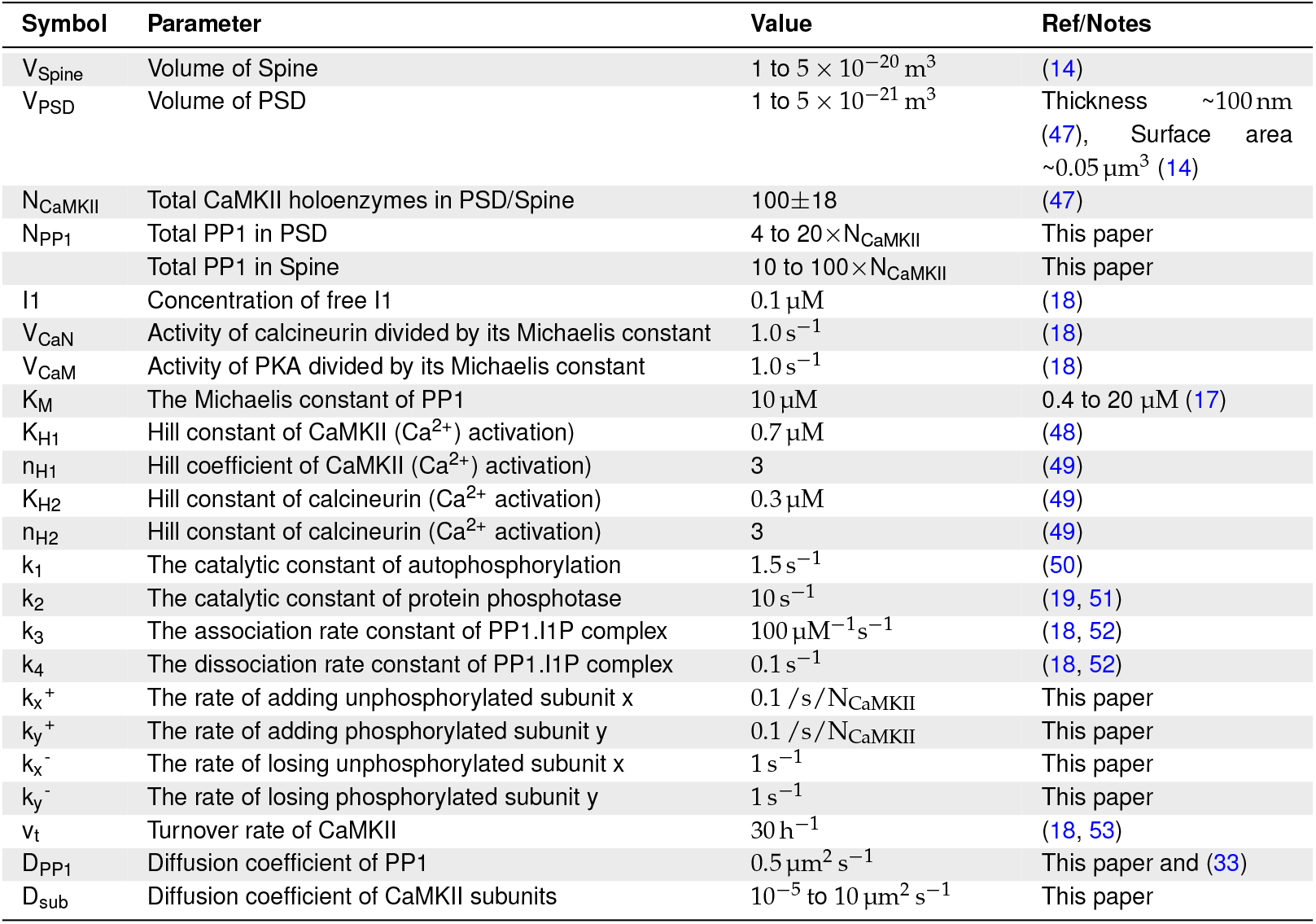
Table of parameters used in model.

## ACKNOWLEDGMENTS

We’d like to thank Marcus Benna, Stefano Fusi and Moitrayee Bhattacharyya for discussions related to their work, Mukund Thattai for useful discussions on the stochastic reaction diffusion methods, and Bhanu Priya for useful comments on the manuscript. This work was funded by NCBS/TIFR and SERB JC Bose fellowship `SB/S2/JCB-023/2016` to `USB`.

**Author contributions**
DS designed the project, carried out the simulations and wrote the paper. USB designed the project and wrote the paper.

